# A mixed blessing of viruses in wastewater treatment plants

**DOI:** 10.1101/2021.11.08.467685

**Authors:** Ling-Dong Shi, Xiyang Dong, Zongbao Liu, Yuchun Yang, Jih-Gaw Lin, Meng Li, Ji-Dong Gu, Li-Zhong Zhu, He-Ping Zhao

## Abstract

Activated sludge of wastewater treatment plants harbors a very high diversity of both microorganisms and viruses, wherein the latter control microbial dynamics and metabolisms by infection and lysis of cells. However, it remains poorly understood how viruses impact the biochemical processes of activated sludge, for example in terms of treatment efficiency and pollutant removal. Using metagenomic and metatranscriptomic deep sequencing, the present study recovered thousands of viral sequences from activated sludge samples of three conventional wastewater treatment plants. Gene-sharing network indicated that most of viruses could not be assigned to known viral genera, implying activated sludge as an underexplored reservoir for new viruses and viral diversity. *in silico* Predictions of virus-host linkages demonstrated that infected microbial hosts, mostly belonging to bacteria, were transcriptionally active and able to hydrolyze polymers including starches, celluloses, and proteins. Some viruses encode auxiliary metabolic genes (AMGs) involved in carbon, nitrogen, and sulfur cycling, and antibiotic resistance genes (ARGs) for resistance to multiple drugs. The former group of virus-encoded genes (i.e. AMGs) may enhance the biodegradation of contaminants like starches and celluloses, suggesting a positive role for viruses in strengthening the performance of activated sludge. However, the latter group (i.e. ARGs) would be disseminated to different microorganisms using viruses as gene shuttles, demonstrating the possibility for viruses to facilitate the spread of antibiotic resistance in the environment. Collectively, this study highlights the mixed blessing of viruses in wastewater treatment plants, and deciphers how they manipulate the biochemical processes in the activated sludge, with implications for both environmental protection and ecosystem security.

## Introduction

Wastewater treatment plants (WWTPs) guarantee both human water security and ecological environment sustainability, by removing noxious pollutants and producing clean water. Activated sludge, since invented more than 100 years ago, has been applied successfully in the modern urban societies and is still the core of current WWTPs ^1^. Microorganisms in the forms of flocs or granules are long believed as the key responsible bio-catalysts of activated sludge, for removing nutrients and micropollutants ^2^. Recent studies suggest viruses, usually many folds more than microbial population, have the potential to impact the activity and functionality of their microbial hosts in various habitats, including ocean ^3^, freshwater ^4^, permafrost ^5^, and other natural ecosystems ^6,7^. However, whether viruses have a contribution in artificial and engineering systems, for example in the activated sludge, and what are their exact roles remain poorly understood.

Owing to a lack of essential genes, prokaryotic viruses are obligate intracellular parasites and rely on microorganisms for their survival and replication ^8^. During the infection cycle, viruses can either lyse microbial cells to release host-derived nutrients, or transduce and express viral genes to reprogram the metabolisms of the infected hosts. One group of such genes is known as auxiliary metabolic genes (AMGs). The first AMG was discovered in marine phages and identified being associated with photosynthesis ^9^. Subsequent bioinformatic mining uncovered more viral AMGs involved in the biogeochemical cycling of carbon ^10,11^, nitrogen ^12,13^, sulfur ^14,15^, and phosphorus ^16,17^. Notably, some viruses also encode genes related to degradation of polysaccharides and were proposed to have contributions to polymer hydrolysis in soil ^5,18^. These studies suggest viruses very likely play a significant role in treatment processes by activated sludge, given the fact that municipal wastewater is dominated by macromolecular organics such as proteins, sugars, and lipids ^19,20^.

In addition to AMGs, another virus-encoded group is antibiotic resistance genes (ARGs), which help microorganisms to resist antibiotics and thus induce worldwide morbidity and mortality ^21^. Currently, about 700,000 people die from the spread of ARGs each year and this number will increase to 10 million by 2050 if no effective intervention is made ^22,23^. ARGs are routinely propagated within and across microbial genera via vertical and horizontal gene transfer (HGT). Viral transduction is one of the most important HGT pathways and has the potential to transfer a large diversity of ARGs locating in viromes ^24-26^. Activated sludge is a hotspot for ARGs and plays critical roles in the selection, proliferation and dissemination of ARGs ^27,28^. Despite a recently reported viral catalogue from activated sludge ^29^, it remains largely unexplored whether viruses expedite the spread of ARGs in those systems.

We hypothesize that viruses can transduce and express viral genes to alter microbial functions and further impact the water quality of effluent discharged after activated sludge treatment. Using *in silico* approaches, we firstly identified and classified viral sequences from bulk metagenomic datasets, then linked the microbial hosts to viruses and clarified their exact contributions, and finally deciphered virus-borne genes potentially associated with wastewater treatments and ecosystem safety. By doing so, we revealed the double-edged roles of viruses in activated sludge being beneficial to removal of both nutrients and pollutants, but detrimental to human health because of ARG spread.

## Results and Discussion

### Viruses in activated sludge are diverse

Three activated sludge samples, designated as Linkou, Bali, and Wenshan, were collected from conventional WWTPs in Taiwan and sequenced deeply to obtain comprehensive profiles of microbial community members (estimated average coverage ≥90%) ^30-33^. Thousands of viral sequences were identified from metagenomic assemblies and clustered at 95% nucleotide identity, finally generating 1830 (Linkou), 2265 (Bali), and 3400 (Wenshan) non-redundant viral operational taxonomic units (vOTUs) at approximately species-level taxonomy ^34^. All the recovered vOTUs had length ≥5 kb, including one longer than 250 kb and possibly belonging to the clade of huge phages. A recent global-scale study reported that huge phages are widely distributed across Earth’s natural ecosystems but are only discovered in one artificial system namely a thiocyanate bioreactor ^6^. Future investigations are warranted to delineate if huge phages are also commonly and actively present in engineering systems like WWTPs.

Together with reference genomes, 2114 recovered vOTUs were clustered into a total of 1518 viral clusters (VCs) at approximately genus level, by comparing shared genes ^35^ (Figure 1a and Table S1). Around 100 VCs were shared between any two of the three activated sludge samples and 33 VCs among all three samples, showing a core viral community similar to bacterial populations (Figure 1b) ^2^. As expected, different sludge samples also possessed their own unique VCs, likely due to the variations of influent wastewater characteristics and the resulting microbial community and hosts ^30^. Interestingly, only a minority of vOTUs derived from activated sludge (∼1.4%) were shared with taxonomically known viruses from NCBI RefSeq (Figure 1b and Table S1), indicating that a large proportion of recovered viruses could not be assigned to known genus-level taxonomy. Among the classified viruses, 18 vOTUs belonged to *Podoviridae*, 11 vOTUs to *Siphoviridae* and 2 vOTUs to *Myoviridae* (Figure 1c), all of which were much lower than the classified viral quantities discovered in permafrost and deep sea sediments ^5,7,18^, but comparable to that in glacier ice ^36^. This suggests that activated sludge harbored diverse and especially the unknown viruses, which may impact the performance of such engineering systems.

**Figure 1.**
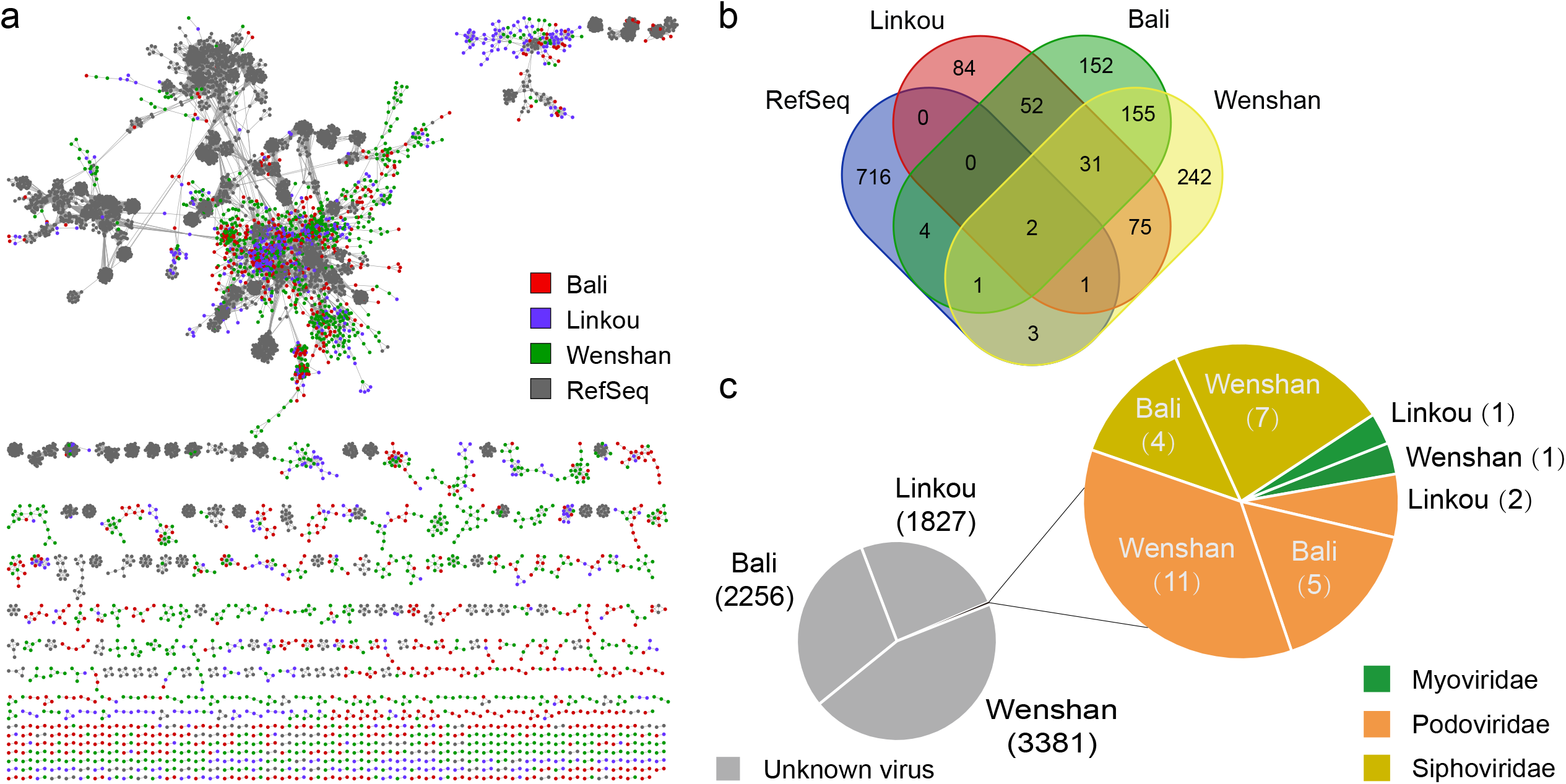
Taxonomic diversity of activated sludge viruses. (a) A gene-sharing network of viral sequences from activated sludge samples in different WWTPs (n=2265 for Bali, n=1830 for Linkou, n=3400 for Wenshan) and RefSeq prokaryotic viral genomes (n=1748). Nodes and edges represent viral genomes and their shared protein content, respectively. (b) Venn diagram indicates shared viral clusters among different activated sludge samples and the RefSeq database. (c) Pie charts indicate taxonomic assignments and relative abundances of vOTUs.

### Viruses infect microbial populations capable of polymer hydrolysis

Viruses play critical roles in habitat biogeochemical processes, one of which is by infecting microbial populations ^37^. Community populations in three activated sludge samples were recovered through assembling and binning, and finally generating a total of 557 metagenome-assembled genomes (MAGs) (173 in Linkou, 183 in Bali, and 201 in Wenshan) after quality filtration (Table S2). To obtain a domain-to-species taxonomy, the Genome Taxonomy Database (GTDB, release 95) was used to classify the recovered MAGs ^38^, possibly leading to different taxonomic assignments for some members from our previous study, for example with the phylum Thermoproteota substituting Thaumarchaeota (Table S2) ^31,33^. Based on CRISPR spacers targeting, genome contents, tRNA sequences, and oligonucleotide frequency patterns, inherent microbial hosts were predicted for a total of 460 vOTUs (137 in Linkou, 137 in Bali, and 186 in Wenshan) (Table S3). These hosts spanned dozens of phyla, with only one archaeal phylum (i.e. Thermoproteota based on the GTDB) found to be infected and the remainder being bacteria (Figure 2a and Table S2). Proteobacteria in the present study were predicted to be infected by the highest number of viruses (150 vOTUs), followed by Bacteroidota by 77 vOTUs, mainly because these members dominated the microbial community of activated sludge (Figure S1).

**Figure 2.**
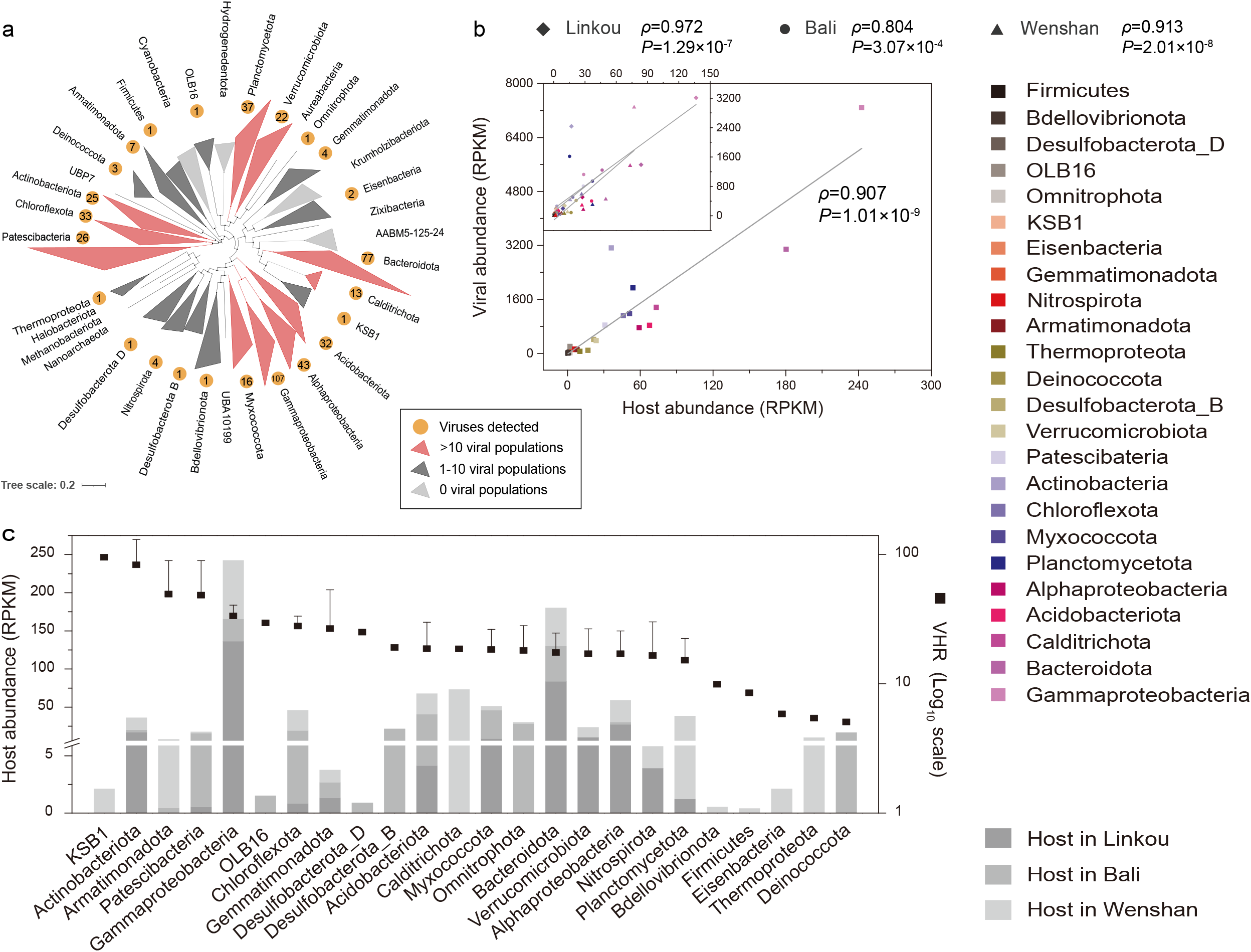
Virus-host linkages and abundance patterns of activated sludge samples. (a) Phylogeny of archaea and bacteria at the phylum level (classes for Proteobacteria) recovered from activated sludge samples. The tree was constructed based on a concatenated set of 122 archaeal or 120 bacterial conserved single-copy marker genes. The lineages indicated by orange circles were predicted to be infected by viruses, with the numbers of vOTUs shown in the circles. (b) Spearman correlation between abundances of viruses and their hosts in individual activated sludge samples (inset) and in combined samples. The colors in symbols indicate different lineages of hosts. (c) Virus/host abundance ratios (VHRs) in activated sludge samples. Color bars represent phylum- or class-level host abundances in different samples. Error bars indicate standard deviation of three samples or mean deviation of two samples.

The predicted virus-host linkages were supported by the comparative abundances of viruses and microbial hosts (Table S3). The replication of viral genomes is dependent on infected prokaryotic cells, that controls viral burst size and thus keeps a typically stable virus-host ratio (VHR) in ecosystems ^39,40^. Viral abundances here showed a significantly positive correlation with host abundances when treating activated sludge samples both individually and collectively (Figure 2b), confirming the accuracy of predicted virus-linked hosts. VHRs ranged from 5.1 to 94.9, with the average value of 26.2 (Figure 2c). By integrating hundreds of publications about viral ecology, mean VHRs were reported between 5.6 and 704.4 in different ecosystems ^40,41^. Activated sludge in the present study had a comparable value to the ocean (i.e. 26.5), which was estimated to contain ∼10^31^ viruses that critically drive the marine biogeochemistry ^42,43^. This indicated a high level of active genome replication of viruses and implied their potential significant roles in activated sludge.

Among the detected viruses, more than half were transcriptionally active, with approximately 10% being linked to microbial hosts (Figure 3). Likewise, transcriptions of dozens of virus-linked hosts were observed. Most of these transcribed and infected microbial populations had high replication rates (iRep values of 1.20-2.20) and were capable of polymer hydrolysis, including starch, cellulose, and proteins that were main constituents of activated sludge ^19,20,44^ (Table S4). Taking the 18 active microbial members from Linkou as an example, 13 MAGs encoded the γ-amylase gene; 13 MAGs possessed the cellobiosidase gene; all of them possessed the acyl-CoA ligase gene; and 15 MAGs contained the endopeptidase gene (Figure 3a and Table S4). These genes are involved in the first step of hydrolysis of the preceding organic substances or pollutants into glucose, cellobiose, and peptide, respectively ^45^. The produced intermediates could be further degraded by various microorganisms to finally decrease chemical oxygen demand (COD). Overall, these results revealed that viruses had an important but previously overlooked contribution to removal of nutrients and pollutants by infecting active microbial populations in activated sludge. On the one hand, viral infection might kill the hosts and thus weaken the biodegradation of pollutants in wastewater. On the other hand, viruses also have abilities to accelerate the overall metabolic reactions of host cells ^46^, for example, by increasing the nitrogen uptake and augmenting energy production ^47,48^, and thus strengthen the treatment efficiency and capacity. At the present, we can neither prove that viruses help to remove nutrients and/or contaminants in activated sludge, nor can we prove that they do not, based on the current available infection information.

**Figure 3.**
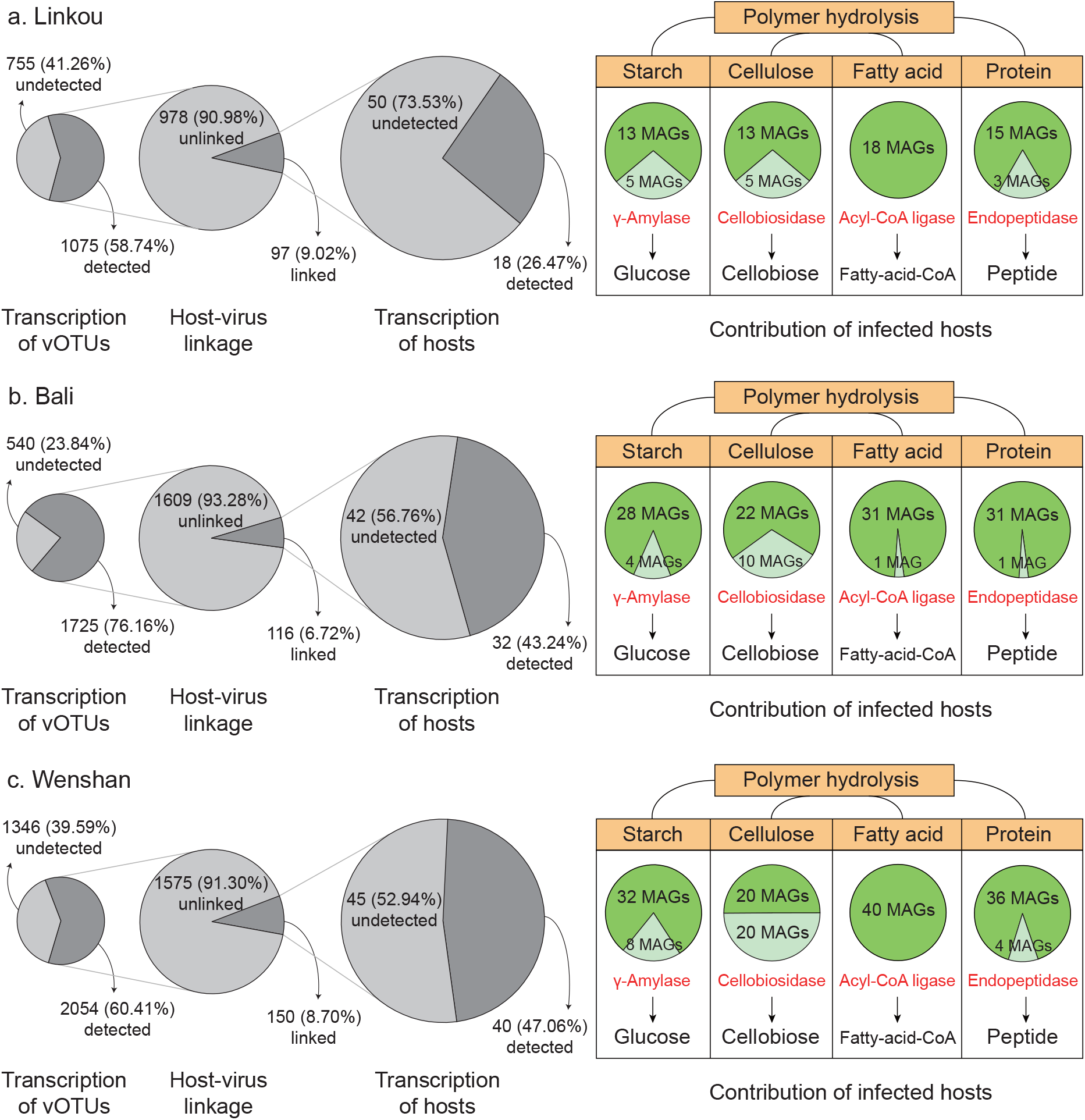
Transcription and functionality of microbial populations infected by viruses. Grey pie charts indicate percentages of transcribed vOTUs, host-linked viruses, and transcribed hosts in samples of Linkou (a), Bali (b), and Wenshan (c). Green pie charts in the right side indicate numbers of transcribed host populations which were predicted to be infected by viruses and harbor genes involved in polymer hydrolysis (dark green). Light green indicates the MAGs with no relevant genes. Key enzymes are highlighted in red.

### Viruses encode AMGs for the removal of pollutants

Encoding and expressing auxiliary metabolic genes is an important pathway for viruses to impact the biogeochemical processes of ecosystems. In activated sludge, a total of 5 AMGs were identified in 21 vOTUs, including genes encoding α-amylase (*amy*), β-glucosidase (*gba*), chitinase (*chi*), nitronate monooxygenase (*nmo*), and phosphoadenosine phosphosulfate reductase (*cysH*) (Figures 4 and S2). All of the AMGs were flanked by viral hallmark genes or viral-like genes, supporting their affiliations to viruses (Table S5) ^49^.

**Figure 4.**
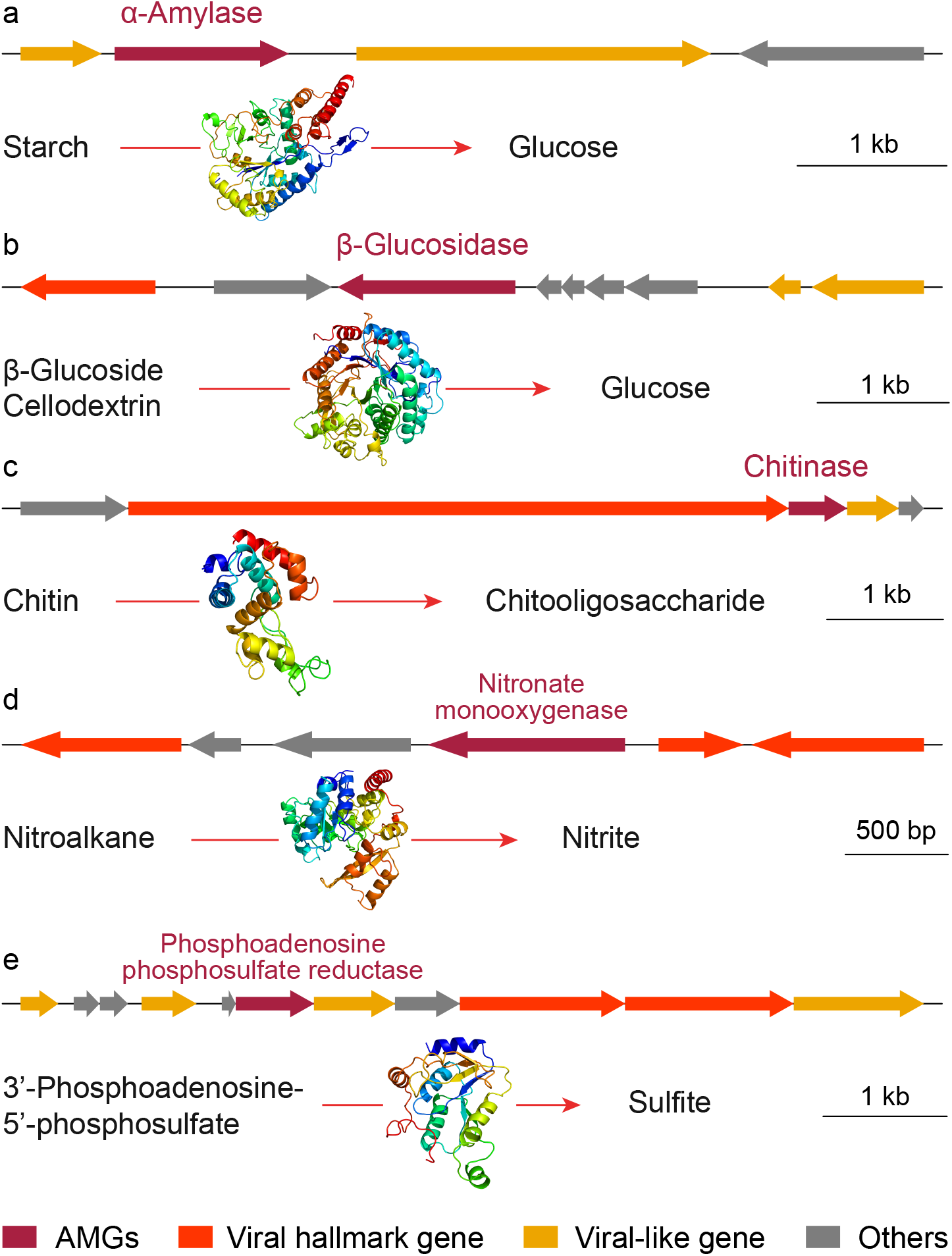
Examples of the adjacent genomic context, predicted functionality, and protein structure of representative viral auxiliary metabolic genes. Dark red arrows indicate the viral AMGs annotated as α-amylase (a), β-glucosidase (b), chitinase (c), nitronate monooxygenase (d), and phosphoadenosine phosphosulfate reductase (e). Viral hallmark and viral-like genes are highlighted in red and orange, and other genes in grey.

The gene *amy* found in a Linkou virus is predicted to randomly cleave α-1,4-glycosidic linkages in starch to generate glucose (Figure 4a) ^50^. The topology of the putative active and catalytic sites of the recovered viral α-amylase conformed to those of functionally known enzymes from microorganisms (Figure S3a). These conserved motifs, together with the tertiary structure from the high-confidence model (Table S6), validated the predicted functions of viral *amy* and highlighted its role in polymer hydrolysis in activate sludge, as important as proposed in peatland soils ^5^. Though the vOTU containing the gene *amy* could not be linked to a identified host, the phylogenetic analysis of α-amylase implied that virus might obtain the AMG from members of Cytophagales during infecting them (Figure S4).

The gene *gba* observed in a Wenshan virus is reported to hydrolyze the β-glucosidic bond between carbohydrate residues for example in β-glucoside and cellobiose with glucose as the product (Figure 4b) ^51^. This protein family was predicted to have no conserved motifs; however, the recovered viral *gba* was modeled to form an identical fold structure to Family 1 glycosyl hydrolase with high confidence (Table S6), suggesting that the β-glucosidase gene, had the genetic potential to degrade complex carbohydrates in activated sludge. The phylogenetic homology between the viral gene and those from *Verrucomicrobium* (Figure S5) was consistent with the predicted viral host belonging to the family Verrucomicrobiaceae (Table S3), both suggesting the origin of viral *gba* being from members of this microbial lineage.

The gene *chi* discovered in viruses from all the three activated sludge samples is capable of cleaving polymer chitin into low-molecular-weight chitooligosaccharide (Figure 4c) ^52^. Viral chitinases possessed conserved catalytic sites and modeling tertiary structures, identical to proteins known to have hydrolytic functions (Figure S3b and Table S6). One of the eight identified vOTUs containing the gene *chi* was linked to a host belonging to Burkholderiaceae (Tables S3 and S5). This was in line with the closest phylogeny of the corresponding AMG to genes from Burkholderiales members, showing the viral infection to this lineage and transfer of chitinase genes (Figure S6). Other viral *chi* genes possibly originated from different microbial populations spanning taxonomic orders, according to the phylogenetic distance. Viral chitinases are routinely to release descendants from infected hosts ^53^. This was supported by the genomic context of vOTU k141_586281, where the chitinase gene comes after all the viral structure genes including those involve in capsid and tail assembly (Table S5). However, some vOTUs like k141_1949521 have chitinase genes locating at the front of the entire contig (Table S5), implying these genes might be expressed along with the viral replication and possibly contribute to the removal of polymer chitin in activated sludge. Biologically hydrolyzing chitin derived from insects and fungi can also provide carbon and nitrogen sources for coexisting microbial members, as observed in marine and soil environments ^54-56^. Further evidences are warranted to test whether viral chitinases have such contributions.

The gene *nmo* encodes nitronate monooxygenase (known as 2-nitropropane dioxygenase previously) and is able to act on nitronate analogues (e.g. nitroalkane) along with the production of nitrate, nitrite and other substrates depending on different substrates ^57^. This gene is identified in viruses for the first time, and was only found in the Bali sample (Figure 4d). The topology of flavin-mononucleotide-binding sites and catalytic motifs in the viral nitronate monooxygenase accorded with those in microbial proteins (Figure S3c). Additionally, the viral *nmo* was modeled to have an identical crystal structure to a putative nitroalkan dioxygenase (Table S6), together proving the genetic potential for degrading nitronate analogues in activated sludge. Based on the constructed phylogeny, the virus was proposed to obtain the *nmo* gene from members of Burkholderiaceae (Figure S7).

The gene *cysH* is involved in reductive assimilation of sulfate and responsible for the reduction of 3’-phosphoadenosine-5’-phosphosulfate (PAPS) into free sulfite ^58^. Though this gene family was predicted to have no conserved motifs, the virus-derived CysH, also named as PAPS reductase and recovered from all three activated sludge samples, formed very similar tertiary structures to functionally known PAPS reductases (Figure 4e and Table S6). This homology suggested that viral *cysH* could enhance the biosynthesis of methionine and cysteine from sulfate assimilation in activated sludge, as well as in other ecosystems ^7,59-62^. Surprisingly, viral *cysH* genes formed discrete clusters from microbial ones and were manually divided into two clades based on the modeling structures (Figure S8 and Table S6). Clade I and II are structurally similar to those purified from bacteria and fungi, respectively (Table S6). This dissimilarity implied that viruses might infect and acquire *cysH* genes from either prokaryotes or eukaryotes.

Taken together, viruses in activated sludge encode AMGs associated with the biochemical cycling of carbon, nitrogen, and sulfur, and with potential implications of pollutant removal. Transcriptions of several viral AMGs were detected (Table S4), suggesting that viruses were expressing these genes and promoting the biodegradation of contaminants. Other viruses encoding AMGs with undetectable transcriptions might not be in the infection cycle when taking the activated sludge samples.

### Viruses encode ARGs for the exacerbation of drug resistance

Besides auxiliary metabolic genes, viruses are known to encode antibiotic resistance genes (ARGs) and transfer them across different taxonomic levels ^63,64^. In total, 14, 24, and 28 viral genes were predicted as ARGs accounting 0.0004%, 0.001%, and 0.0007% of all the genes in Linkou, Bali, and Wenshan, respectively (Table S7), which were comparable to those reported in other metagenomic studies ^65^. Virus-associated ARGs belonged to various families and conferred resistance prevailingly against elfamycin, aminoglycoside, phenicol, and multiple drugs in all three activated sludge samples (Figure 5a and Table S7). Except for the former one, aminoglycosides and phenicols have been widely used in clinical antibacterial chemotherapy. Aminoglycosides are potent and broad-spectrum antibiotics which protect humans by inhibiting the protein synthesis of pathogens ^66^. This class includes streptomycin, kanamycin, and gentamicin, and is always effective for many microbial infections. Phenicols against both Gram-positive and -negative bacteria can destroy their protein synthesis by blocking the peptide elongation ^67^. Chloramphenicol is the representative in this bacteriostatic antibiotic class, and mainly used to treat eye and ear infections. Because of the widespread applications of such antibiotics, their residues and corresponding resistance genes are both frequently detected in natural estuaries, swine manure, and urban sewage ^68-71^. Given the co-occurrence and transcription of viruses and virus-borne ARGs in activated sludge (Table S7), viral transduction may contribute to the global increase of antibiotic resistance.

**Figure 5.**
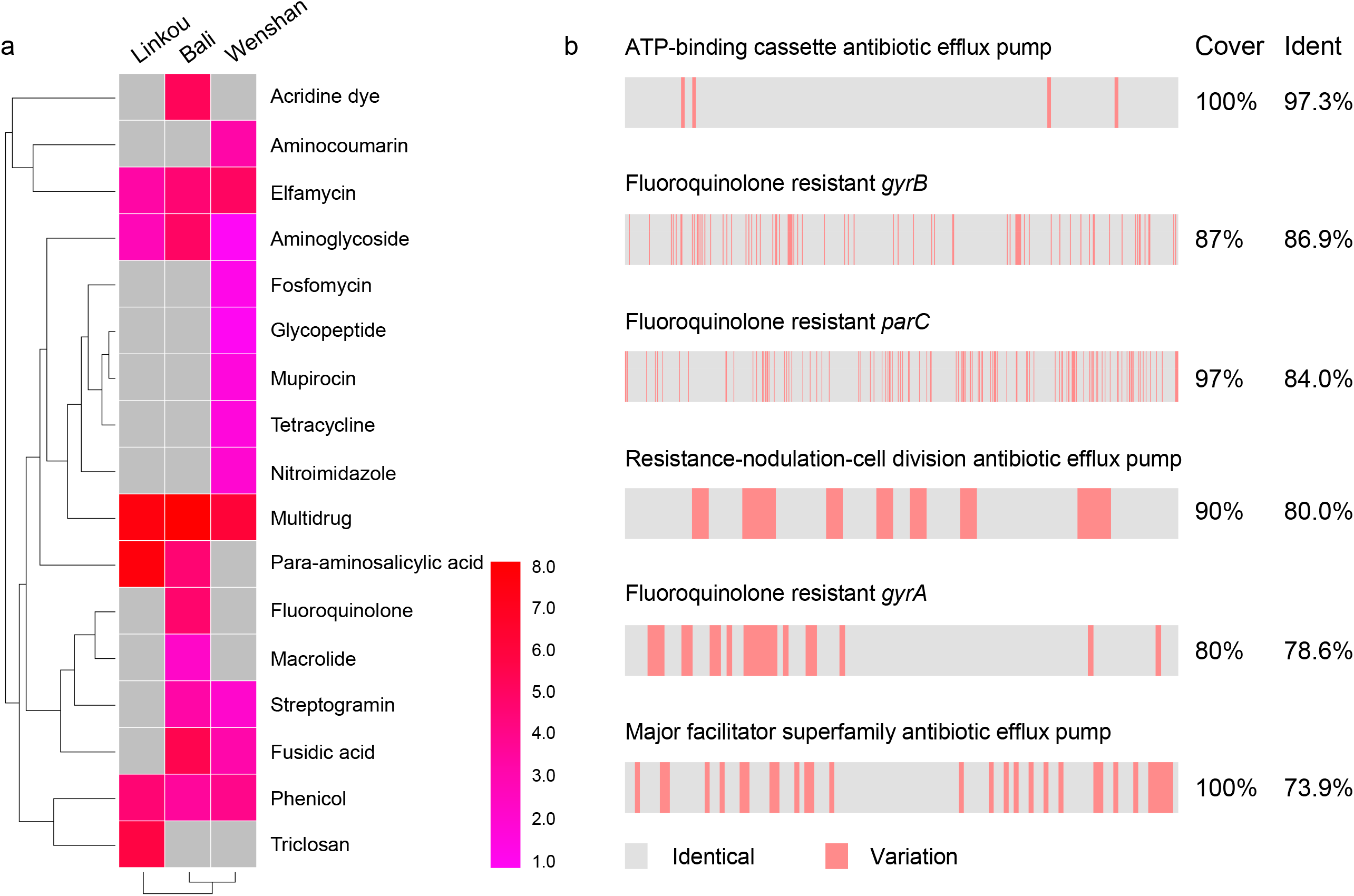
Resistant drugs and sequence alignments of selected viral antibiotic resistance genes. The abundances of viral ARGs in heatmap (a) were normalized by a logarithm base of 2. Grey squares indicate the absence of corresponding genes in the sample. Grey and red regions in (b) indicate identical and variable nucleotide sequence sites, respectively, in relative positions of genes from viruses and their hosts. The coverage and identity were calculated by the BLASTn algorithm.

To evaluate such role of viruses in activated sludge, ARGs encoded by host-linked vOTUs were selected and compared to the genomes of microbial hosts. Homologs of all the selected viral ARGs could be found in the corresponding host genomes, with the nucleotide identity ≥70.0% and coverage ≥80.0%, including genes conferring resistance to fluoroquinolones and multiple drugs (Figures 5b and S9). Notably, an ARG recovered from the Bali sample and annotated as ATP-binding cassette antibiotic efflux pump had a small variation by only four amino acid residues from its host-derived homolog (Figure S9). This pump uses the energy from ATP binding/hydrolysis and transports antibiotics out of cells, and thus conferred resistance on living organisms ^72^. Overall, ARGs carried by viruses and similar to hosts indicated that viruses very likely transduced and propagated ARGs between microbial populations and finally exacerbated the drug resistance in ecosystems. Even worse, some viruses could infect pathogenic microorganisms – for example the two Bali vOTUs predicted to infect Burkholderiaceae members that contain pathogens (Tables S3 and S7) ^73^ – and thus likely pose more severe threats to human health.

Antibiotic resistance has become a global health crisis and may lead to one death every 3 s by 2050 ^22^. Horizontal gene transfer (HGT) via conjugation, transformation, and transduction is the primary cause of the dawn. Compared to the former two pathways having been studied in activated sludge ^74,75^, the latter one i.e. transduction is largely unexplored. Consistent with a previous study reporting rare ARGs encoded by phages ^76^, our study only identified 14-28 candidate genes of recovery frequencies at 4×10^−6^∼1×10^−5^, using more conservative thresholds with much lower e-values and higher identity and bit-scores (Table S7). Particularly, most of the identified ARGs are not associated with antibiotic efflux, which would reduce the over-estimation induced by the CARD category “drug transporters” ^76^. Among the ARG-carried viruses, ∼10% were predicted to have lysogenic cycles based on the presence of integrases (Table S7). However, they can turn into lytic styles triggered by very common environmental conditions in WWTPs including solar radiation and shock from chemical pollutants ^77,78^, making it not impossible to transfer genes between microbial hosts. On the basis of such conservative explorations, we therefore propose that viral transduction has a potential contribution to ARG spread, which helps to partially explain the enrichment of ARGs in activated sludge ^27^. Significantly, some ARG-carried viruses are predicted to infect pathogenic bacteria, not only found in conventional WWTPs in this study but also in hospital fecal and wastewater samples ^79-81^. Although the impact of HGT on disease frequency remains currently unknown, viruses serving as an ARG shuttle are closely related to human health and should be seriously attended. Had pathogens concentrated multiple antibiotic resistances together, one may have no cure or effective measure when being infected ^82^.

### Perspectives

Because of the pathogenicity of emerging and diverse human viruses ^83,84^, the traditional view has long believed that viruses in activated sludge were pernicious only and should be removed thoroughly. The present study highlighted the mixed blessing of prokaryotic viruses in activated sludge being beneficial to microbial removal of pollutants and detrimental to human health on account of ARG spread (Figure 6). By infecting active microbial populations and expressing auxiliary metabolic genes, viruses could potentially regulate the biochemical reactions of activated sludge and promote the biological hydrolysis of polymers. This would finally help to decrease COD from wastewaters and facilitate the effluent meeting the discharge standard. For another, antibiotic resistance genes are alternative candidates that can also be transferred across different microbial hosts by viruses. This study provided evidences for the virus-mediated potential transduction of ARGs therefore very likely to exacerbate the drug resistance and threat human health (Figure 6). Future studies are needed to isolate representative viruses and their microbial hosts to experimentally decipher and validate their interactions and activities, as well as disclose spatial and temporal patterns of viral infections and contributions given the large dissimilarity of host profiles among different activated sludge samples (Figure S10). Collectively, this study challenges the traditional view of viral activities in activated sludge, and sheds new light on the underexplored but significant functionality of viruses that is necessary for developing next-generation technology of wastewater treatment.

**Figure 6.**
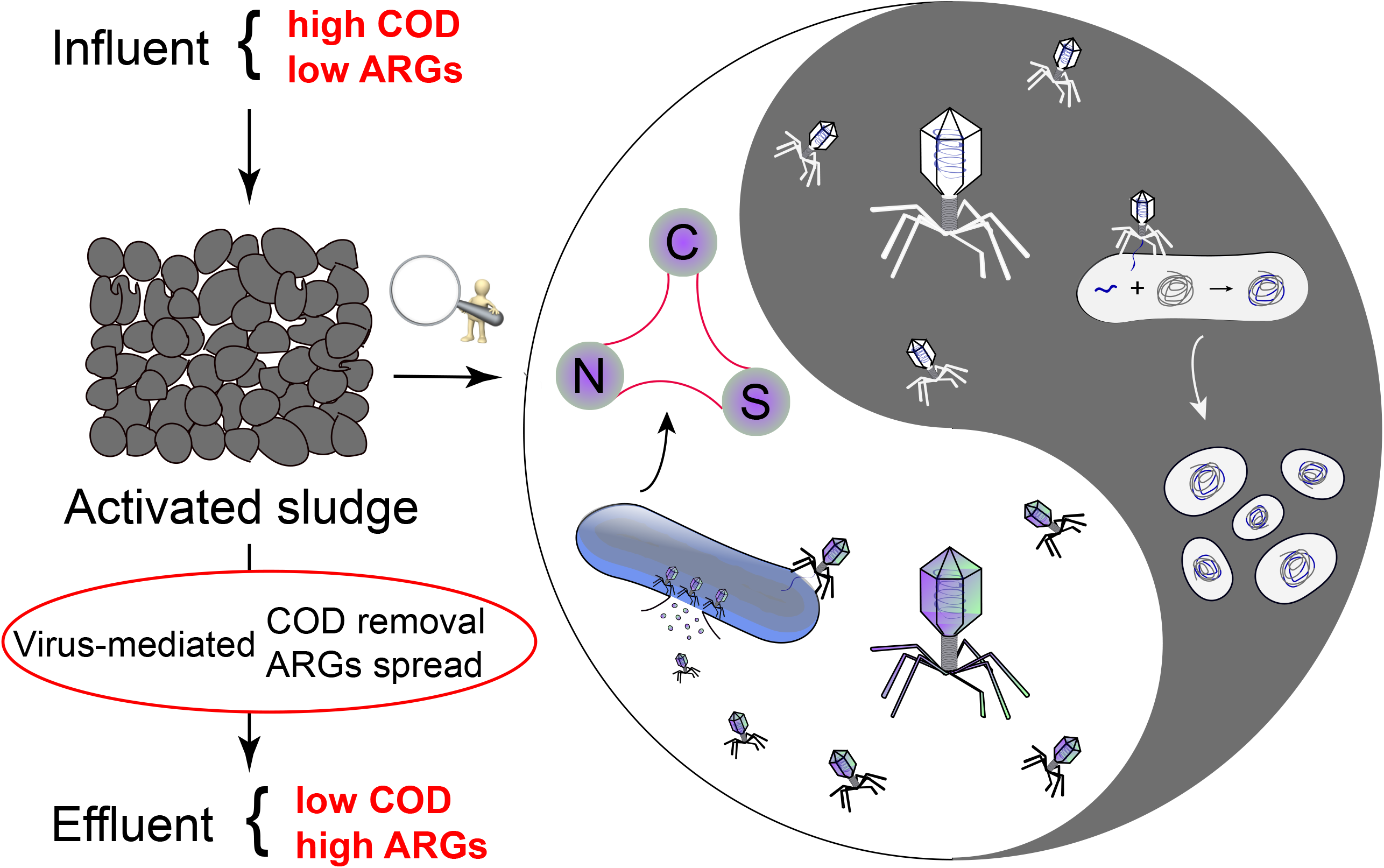
Conception of the double-edged roles of viruses in activated sludge. Activated sludge removes COD but increases the distribution of ARGs from influent to effluent. White panel in the right circle indicates virus-mediated COD removal, and grey panel indicates ARGs dissemination transduced by viruses.

## Materials and Methods

### Collection of sequencing datasets from activated sludge in WWTPs

Metagenomic and metatranscriptomic sequencing reads of three activated sludge samples were downloaded from the NCBI Sequence Read Archive with accession number PRJNA406858 ^30^. Those samples were collected from three conventional WWTPs located in Taiwan, namely Linkou, Bali, and Wenshan. Total DNA and RNA were extracted from the precipitates after centrifuging, which might lose viruses in some extent ^85^. In total, approximately 227.3-258.6 million paired-end reads were obtained for metagenomic analyses and 172.0-182.4 million paired-end reads for metatranscriptomic analyses. Average coverage of metagenomic datasets was estimated at 91.2%, 94.9%, and 90.9% by Nonpareil 3 ^86^. Detailed information of sampling and sequencing is described in our previous study ^30^.

### Metagenomic sequence processing and assembly

Metagenomic raw data was first processed by trimming or removing low-quality reads using Trimmomatic v0.38 with the default parameters ^87^. The quality of filtered reads was evaluated by FastQC v0.11.8 (http://www.bioinformatics.babraham.ac.uk/projects/fastqc/). Then MEGAHIT v1.2.7 was used to assemble reads into contigs for each sample with the kmer set of 21, 29, 39, 59, 79, 99, 119, 141 ^88^. The contigs shorter than 500 bp were excluded from downstream analyses.

### Recovery and annotation of microbial population genomes

Contigs from each assembly were binned into species-level metagenome assembled genomes (MAGs) using MetaBAT2 v2.12.1 with default parameters except for –m 1500 ^89^. Quality of MAGs were assessed by CheckM v1.0.18 ^90^, and MAGs with completeness <50% or contamination >10% were discarded. Taxonomy of MAGs was initially assigned by the GTDB-Tk toolkit v1.3.0 using the Genome Taxonomy Database (release 95) ^38,91^. The assignments were validated by phylogenetic trees that were inferred from a concatenated set of 122 archaeal or 120 bacterial conserved single-copy marker genes and constructed using FastTree v2.1.11 with the WAG+GAMMA models ^92^. Replication rates of microbial populations were estimated by iRep v1.10 ^44^. Gene and protein sequences in each MAG were predicted and annotated by MetaErg v1.2.0 ^93^.

### Identification of viral sequences

Viral sequences were recovered from the preceding metagenomic assemblies. Contigs with length ≥5 kb were identified as viral following the criteria below ^94^: (1) VirSorter2 v2.1 score ≥0.9 ^95^; (2) VirFinder v1.1 score ≥0.9 along with p<0.05 ^96^; (3) both VirSorter2 score ≥0.5 and VirFinder ≥0.7, p<0.05; (4) both VirSorter2 score ≥0.5 and VIBRANT v1.2.1 validation ^97^. The identified viral contigs were then clustered by CD-HIT v4.8.1 at 95% nucleotide identity using –c 0.95, -aS 0.8, and other default parameters ^98^, generating 1830, 2265, and 3400 viral operational taxonomic units (vOTUs) for Linkou, Bali, and Wenshan samples, respectively. The quality of recovered viral populations was assessed by CheckV v0.7.0 ^99^. Life styles were inferred by VIBRANT and CheckV, and considered “lysogenic” for viruses if provirus integrase genes were identified. Otherwise, they were considered “undetermined”.

### Taxonomic assignment of viral contigs

To assign the taxonomy of vOTUs, viral genomes in the NCBI Reference Sequence (RefSeq) Database (release 205) were downloaded as references, and protein sequences of all viral genomes were predicted using Prodigal v2.6.3 with the “meta” mode ^100^. Then the whole genome gene-sharing profiles were built by vConTACT2 v0.9.22 according to the published protocol ^35^, followed by the network visualization using Cytoscape v3.8.0 ^101^. Genus- or family-level taxonomy was assigned to the viral genomes recovered from activated sludge, if they were located in the same viral cluster (VC) with reference genomes. For those had no reference genomes in the same VC or VC subcluster, their taxonomies were temporarily classified as “unknown”.

### Prediction of virus-host linkages

In general, viral populations were linked to microbial members based on: (1) similarities of spacer sequences in viral genomes and microbial clustered regularly interspaced short palindromic repeat (CRISPR) regions, (2) shared genomic contents, (3) homologies of transfer RNA (tRNA) sequences, and (4) similarities in oligonucleotide frequency patterns. Microbial CRISPR arrays in (1) were predicted from metagenomic assemblies by MetaErg. Spacer sequences were parsed according to the repeat regions. Bowtie v1.2.3 was then used to match spacers against viral contigs with ≤1 mismatch ^102^. Once matched, the CRISPR arrays were parsed and assigned to the prokaryotic MAGs, thus linking viruses to microbial hosts. Neighboring protein-coding motifs with distance <10 kbps from matched CRISPR regions were annotated ^103,104^, and only hits with *cas* genes locating adjacently were considered to be confident.

Shared genomic contents in (2) were determined by comparing vOTUs sequences to microbial MAGs using BLASTn v2.9.0 ^105^, with nucleotide identity ≥70%, bit score ≥50, e-value ≤0.001, and alignment length ≥2,500 bp. The tRNA sequences in (3) were annotated from metagenomic assemblies by MetaErg as well, and then parsed and assigned to viral contigs or prokaryotic MAGs. Identified viral tRNAs were compared to microbial ones using BLASTn, with nucleotide identity and sequence coverage both ≥90%. To compare oligonucleotide frequency patterns between vOTUs and MAGs in (4), VirHostMatcher v1.0.0 was performed using default parameters, with d_2_^*^ values ≤0.2 as a positive linkage ^106^.

If multiple and different hosts were predicted for one vOTU, the single host was selected following the priority order below ^5^: (1) CRISPR linkage with adjacent *cas* gene; (2) shared genomic contents or tRNA homology; (3) oligonucleotide frequency match. The virus-host linkage meeting the highest ranking standard was finally chosen.

### Abundance profile of viruses and hosts

Relative abundances of viruses and microbial hosts were calculated in the unit of Reads Per Kilobase per Million mapped reads (RPKM). For viruses, reads after quality control were first mapped to viral contigs using “make” command in CoverM v0.6.1, to make BAM files (https://github.com/wwood/CoverM). Then “filter” command was used to remove the low-quality alignments with reads identity ≤95% and aligned percent ≤75%. Finally the “contig” command was run to calculate the coverage for individual viral contigs using the qualified alignments with the same cutoff parameters in the preceding, namely minimum reads identity of 95% and minimum aligned percent of 75%. For hosts, the calculation protocols were basically same as those for viruses, except for the last step using the “genome” command for microbial MAGs instead of “contig”. Spearman correlations between viral abundance and host abundance were analyzed using IBM SPSS Statistics for Windows v22.0 (IBM Corp., Armonk, N.Y., USA) for separate activated sludge samples and combined samples. Total abundances of infected microbial populations were calculated by summing up all the hosts in the same taxonomy.

### Metatranscriptomic sequence processing and mapping

Metatranscriptomic reads were first quality-controlled by Trimmomatic with the default parameters ^87^, and were then evaluated by FastQC. Ribosome RNA (rRNA) sequences were removed using SortMeRNA v2.1 based on the SILVA and RFAM reference databases ^107^, generating datasets mainly consisted of messenger RNA (mRNA) sequences. Transcriptions were calculated for viruses and hosts using CoverM, following the same protocol as described in the last section, and only those with values above 0 were considered as transcriptionally active.

### Identification of auxiliary metabolic genes and antibiotic resistance genes

Identification of AMGs was based on VIBRANT and DRAM v1.2.0 ^49,97^, after removal of host-derived contamination by CheckV. For VIBRANT, identified viral genomes were directly annotated using “virome” mode; for DRAM, VirSorter2 was first performed to predict gene sequences and generate a contig affiliation file for viral genomes, and then the prepared files were passed to DRAM for the annotation using the viral mode. Annotated AMGs were manually verified based on the genomic contexts ^108^. Conserved domains of identified AMGs were searched by NCBI tool ^109^. Protein structures were modelled using PHYRE2 v2.0 to confirm and resolve the functional predictions ^110^. Phylogenetic trees were constructed using FastTree with the WAG+GAMMA models ^92^ after aligning sequences by MUSCLE v3.8.1551 ^111^. Only branches with ≥50% bootstrap support were shown.

To identify ARGs, gene sequences predicted from viral genomes were searched against the Comprehensive Antibiotic Resistance Database (CARD, v3.1.1) using DIAMOND v0.9.25.126 ^112-114^, with e-value ≤10^−5^ and identity ≥50%. Predicted viral ARGs were compared to their host MAGs to find a sequence homology, using BLASTn with nucleotide identity ≥70% and sequence coverage ≥80%. Relative abundances were calculated in the unit of RPKM using the script in CoverM.

## Data availability

The viral sequences and metagenome assembled genomes recovered in this study have been deposited into the figshare at https://figshare.com/s/61e4a602f161c04b55e8. All other data are available from the corresponding author upon request.

## Acknowledgments

The authors greatly thank the “National Natural Science Foundation of China (Grant No. 51878596, 32061133002)”, the “Key Research and Development Program of Zhejiang Province (Grant No. 2021C03171, 2020C03011)”, the “National Key Technology R&D Program (2018YFC1802203)”, and the “Science and Technology Projects in Guangzhou (No. 202102020970)” for their financial support. They also very much thank Lin-Xing Chen and Donald Pan for the helpful discussion and the constructive suggestions for this work. The author Ling-Dong Shi would like to thank the support from Shanghai Tongji Gao Tingyao Environmental Science & Technology Development Foundation.

## Competing interests

The authors declare no competing interests.

## Notes

### Competing Interest Statement

The authors have declared no competing interest.

