## Supplemental figures for "A mixed blessing of viruses in wastewater treatment plants"

**This PDF file includes:**

Figures S1 to S10.

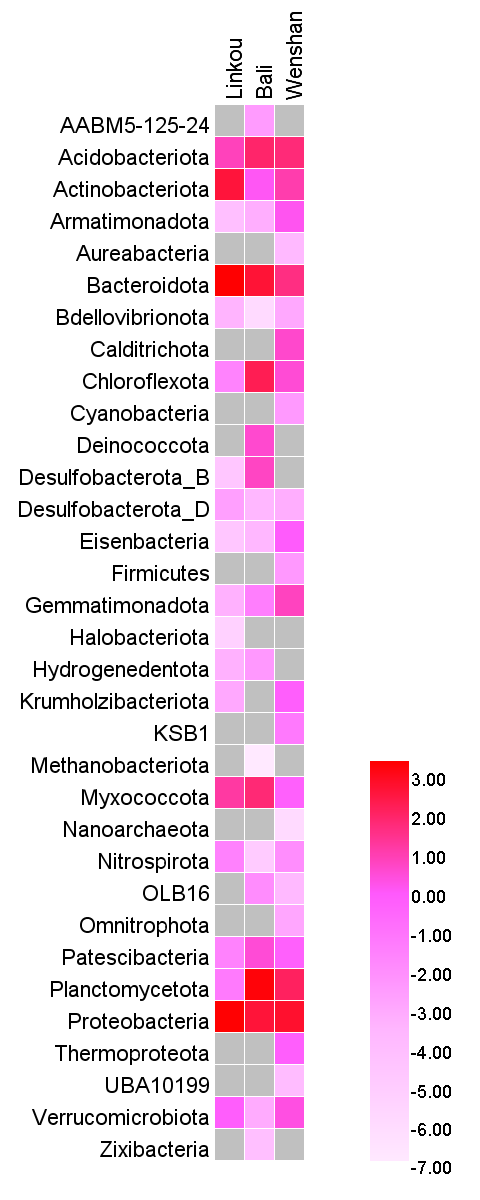

**Figure S1. Relative abundances of microbial phyla in activated sludge.** The percentage values were normalized by a logarithm base of 2. Grey squares indicate the absence of populations from this group.

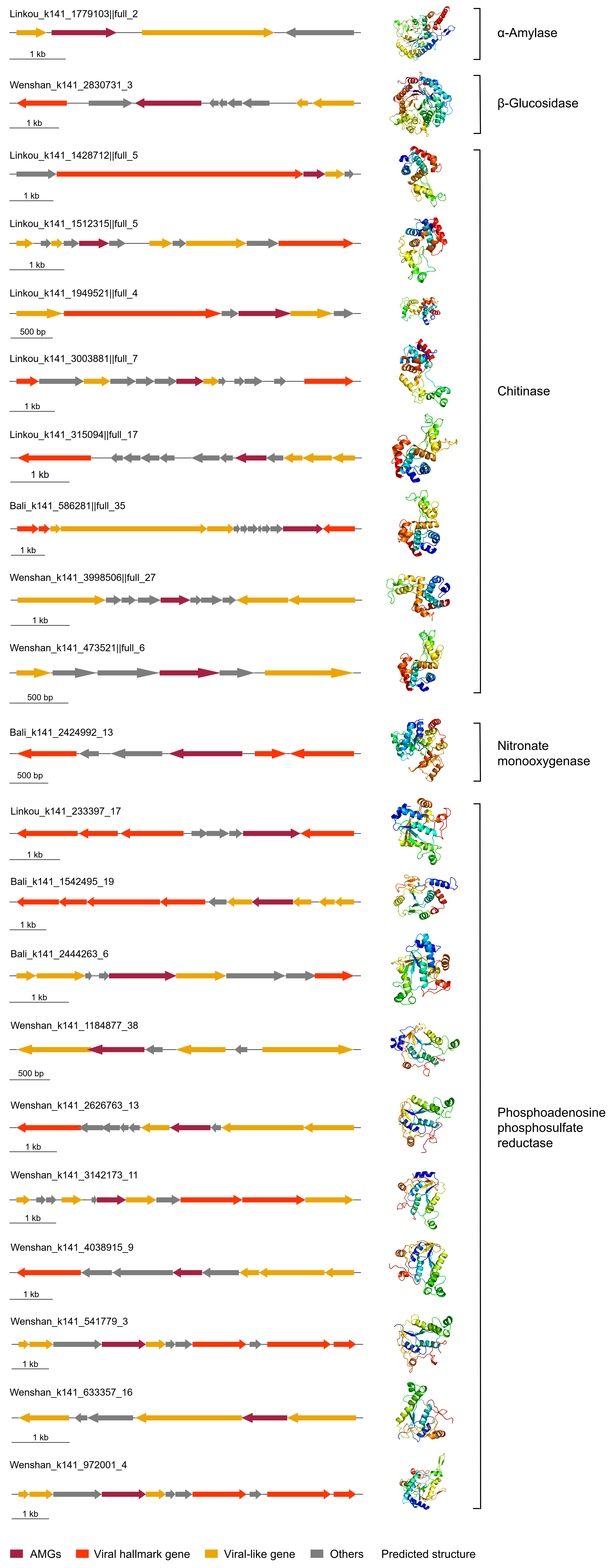

**Figure S2. Adjacent genomic context and protein structure of viral auxiliary metabolic genes.** Titles of the gene operons begin with the sampling fields (i.e. Linkou, Bali, and Wenshan). Annotated AMGs, viral hallmark genes, viral-like genes, and other genes were highlighted in dark red, light red, orange, and grey, respectively.

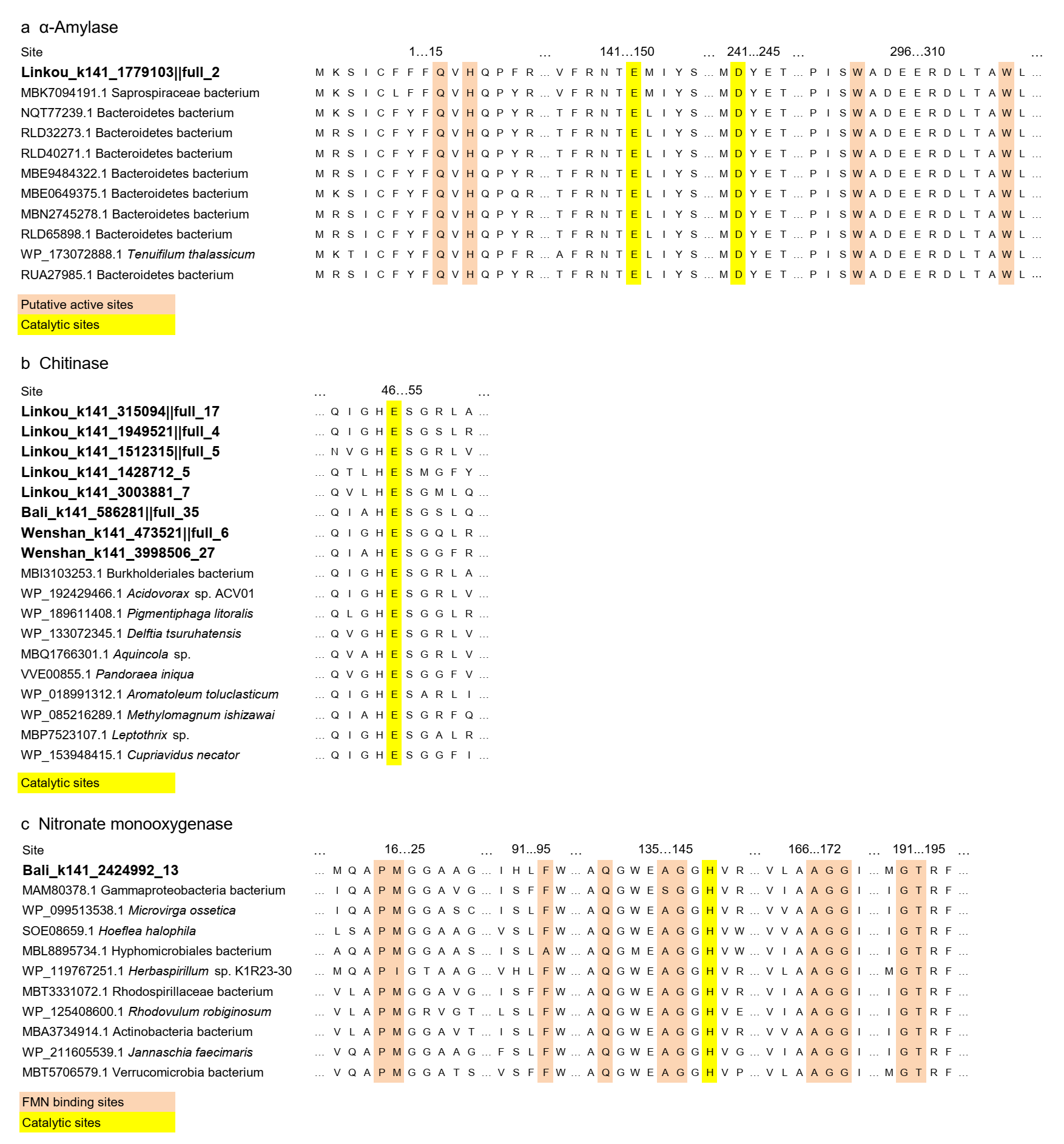

**Figure S3. Alignment of viral and bacterial protein sequences.** Sequences were annotated as α-amylase (a), chitinase (b), and nitronate monooxygenase (c), with conserved catalytic or binding sites highlighted. The names of virus-associated sequences were in bold.

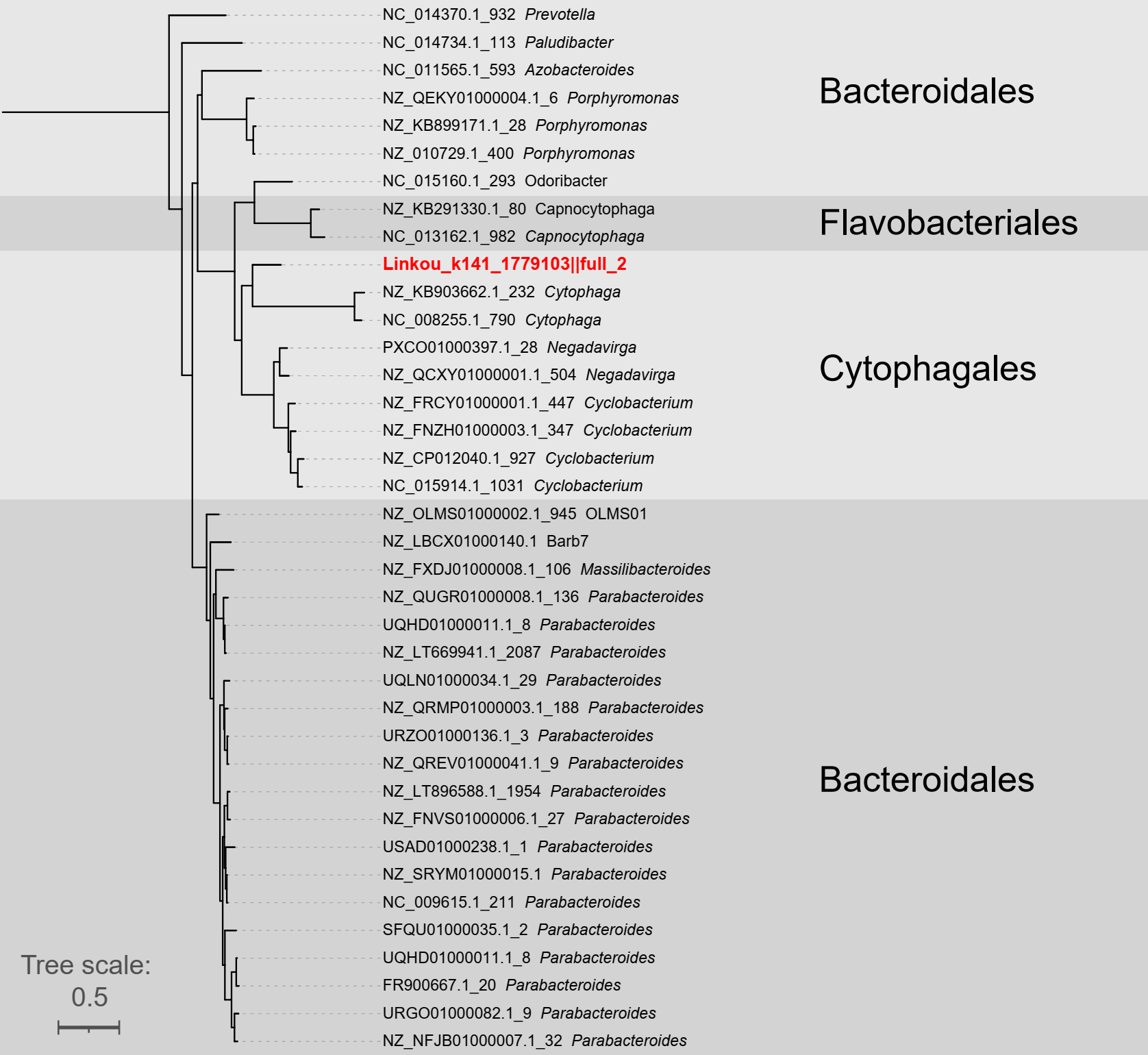

**Figure S4. Phylogeny of α-amylases.** The only viral sequence was highlighted in red and bold. Taxonomy was assigned at the order level, based on the GTDB database.

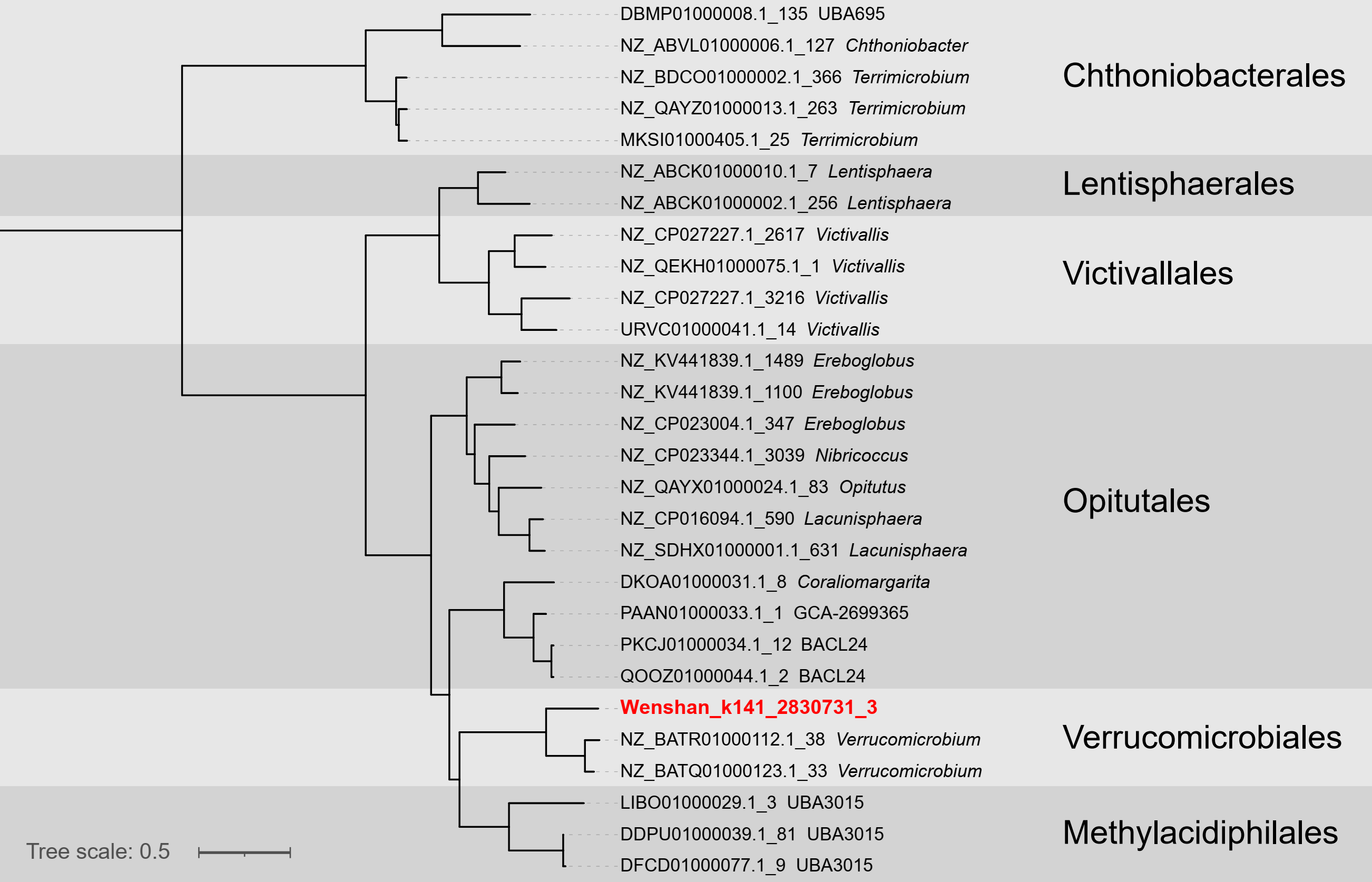

**Figure S5. Phylogeny of β-glucosidases.** The only viral sequence was highlighted in red and bold. Taxonomy was assigned at the order level, based on the GTDB database.

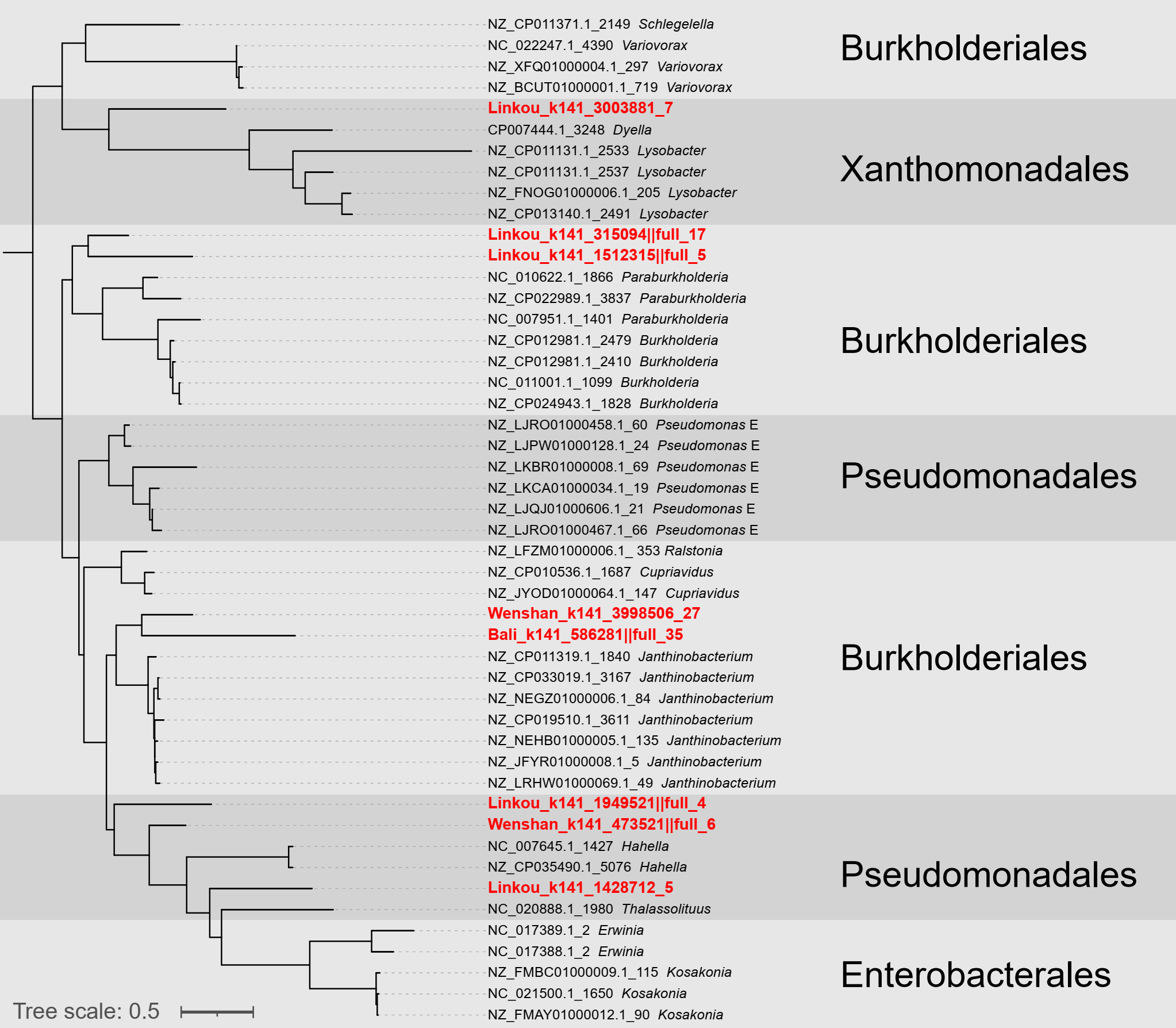

**Figure S6. Phylogeny of chitinases.** Viral sequences were highlighted in red and bold. Taxonomy was assigned at the order level, based on the GTDB database.

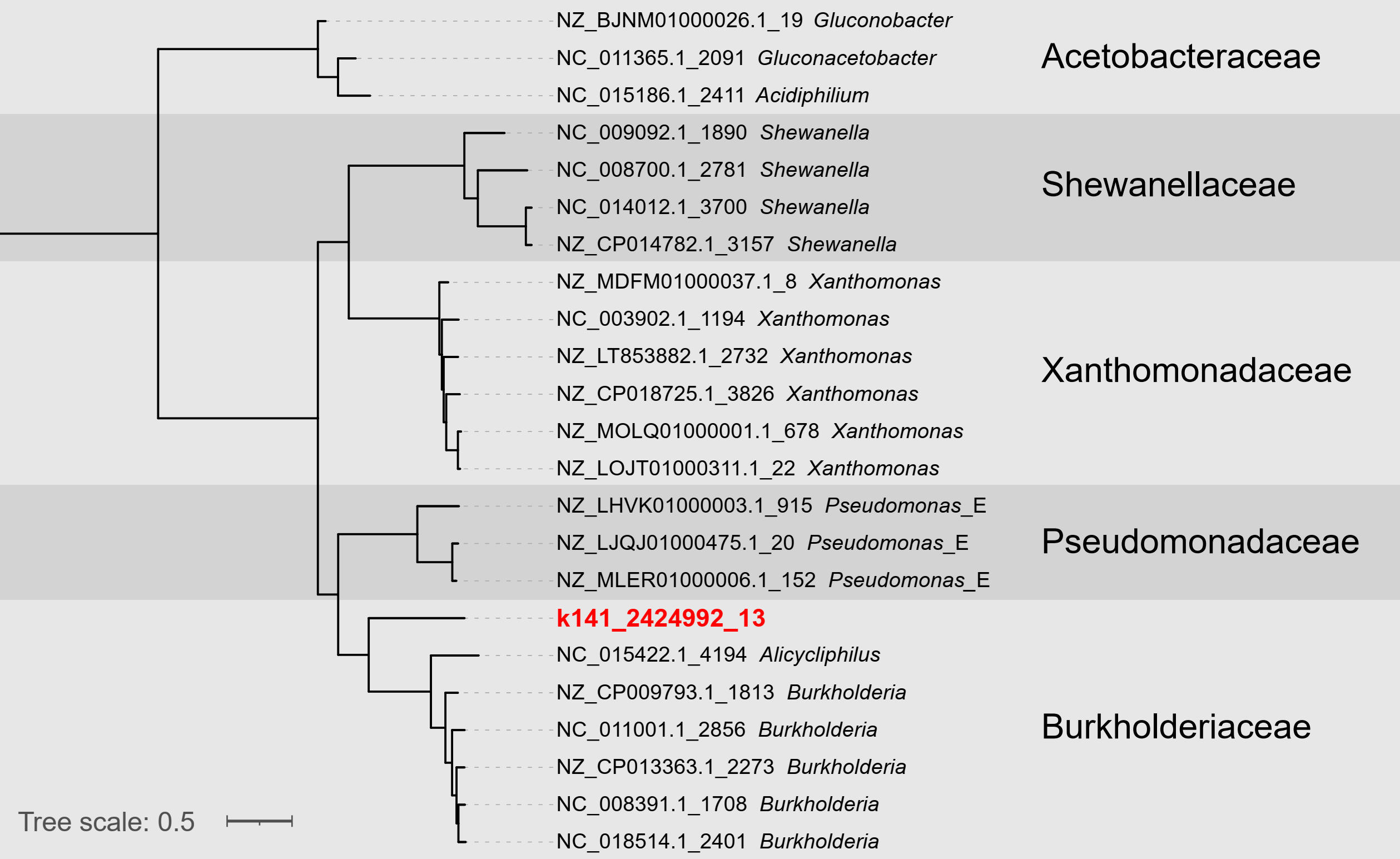

**Figure S7. Phylogeny of nitronate monooxygenases.** The only viral sequence was highlighted in red and bold. Taxonomy was assigned at the family level, based on the GTDB database.

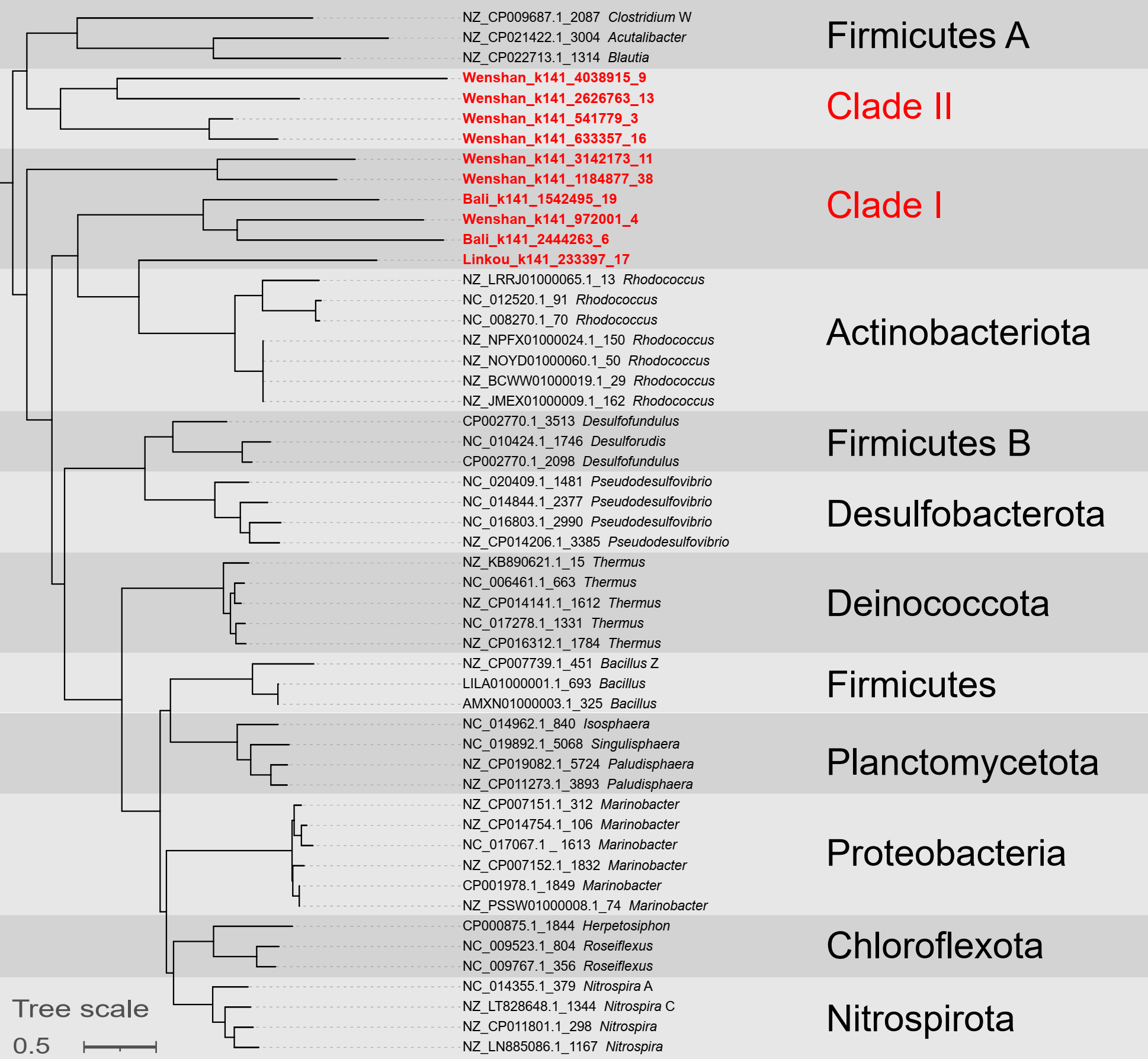

**Figure S8. Phylogeny of phosphoadenosine phosphosulfate reductases.** Viral sequences were highlighted in red and bold. The Clade I proteins have similar fold structures to prokaryotic reductases while Clade II to yeast reductases as modeled (Table S6). Taxonomy was assigned at the phylum level, based on the GTDB database.

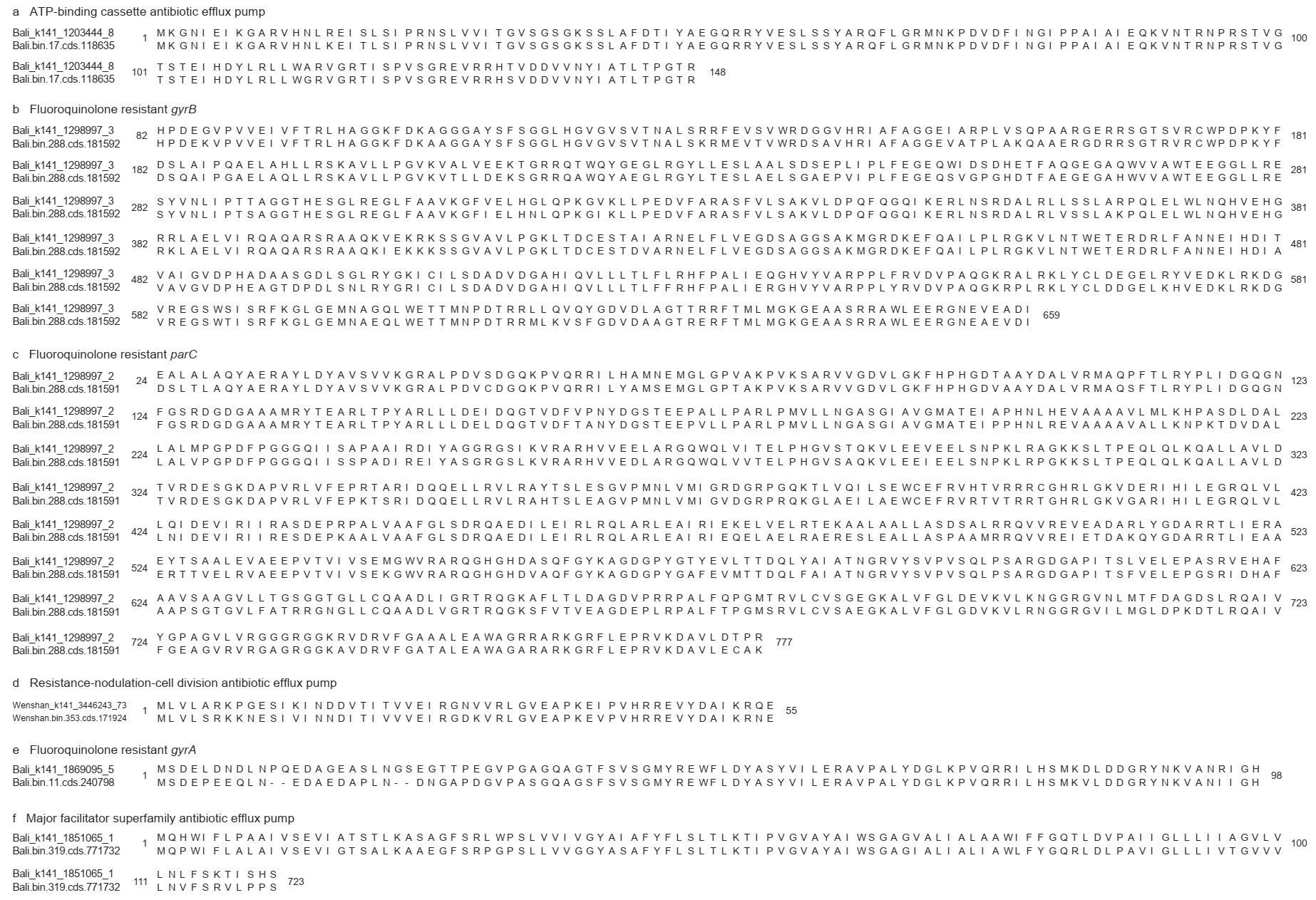

**Figure S9. Alignment of protein sequences recovered from viruses and their hosts.** Sequences were annotated as ATP-binding cassette antibiotic efflux pump (a), fluoroquinolone resistant GyrB (b), fluoroquinolone resistant ParC (c), resistance-nodulation-cell division antibiotic efflux pump (d), fluoroquinolone resistant GyrA (e), and major facilitator superfamily antibiotic efflux pump (f).

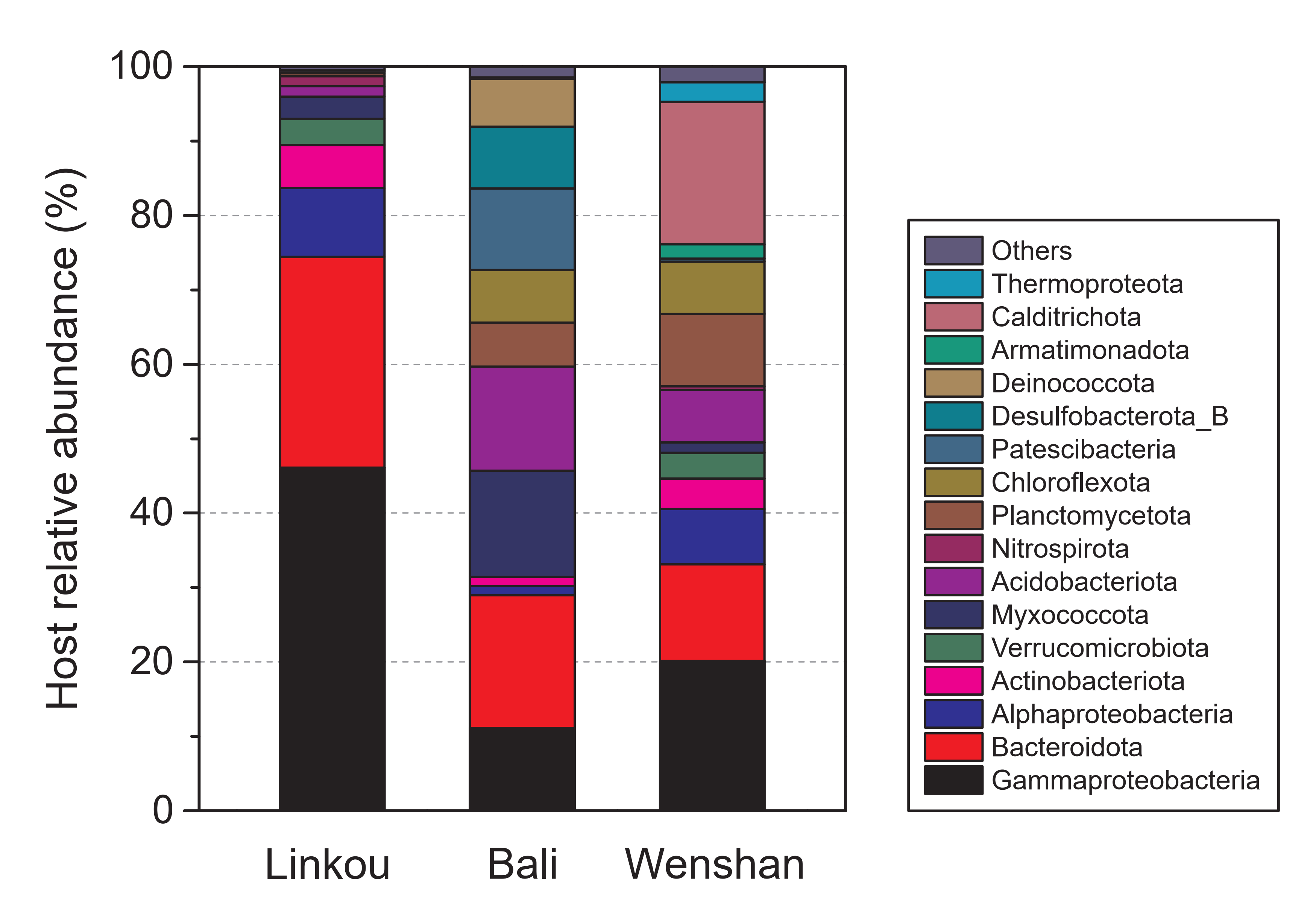

**Figure** **S10. Relative abundances of microbial populations among infected hosts.** Taxonomy was assigned to the phylum level (classes for Proteobacteria), based on the GTDB database. “Others” indicate the sum of populations of which relative abundances were below 1.0%.
